# WARMING AND INCREASED RAINFALL REDUCE BUMBLEBEE QUEEN FITNESS

**DOI:** 10.1101/2025.01.09.632115

**Authors:** C. Ruth Archer, Paul Schmid-Hempel, Regula Schmid-Hempel, Lena Wilfert

**Affiliations:** Institute for Evolutionary Ecology and Conservation Genomics, University of Ulm, Ulm, Germany; Institute of Integrative Biology (IBZ), ETH Zürich, Zürich, Switzerland

**Keywords:** Bumblebee, climatic change, demography, parasites, pollinators

## Abstract

1. How populations respond to climatic change depends on how climate affects individual growth, survival and reproduction – traits that ultimately shape population dynamics. However, climate can have complex effects on these traits and long-term data linking insect life-histories and climatic variation are rare.
2. Here, we test how climate affects health, survival and reproduction in wild *Bombus terrestris* queens caught over 15 years after emerging from hibernation. We relate climate variation experienced by queens as they developed in their natal colonies and entered hibernation, to fitness traits assayed under constant lab conditions.
3. We show that wet years consistently reduced queen fitness, while warm temperatures had positive and negative impacts. Behind these annual effects, lies strong seasonality. In particular, climatic conditions experienced by young queens as they forage, mate and enter hibernation are vital in determining whether they reproduce the following spring.
4. Results suggest that these climate effects are partly due to resources: climatic drivers that reduce resource acquisition prior to hibernation, or disrupt diapause and thus accelerate resource loss over winter, are especially detrimental to spring queen fitness. While results show potential for increasingly negative effects of climate change on bumblebees, they also inform mitigation strategies; providing high-quality forage in late summer, before queens enter diapause.

## INTRODUCTION

Climate change is having costly effects on biodiversity and, as the climate changes further, extinction risk is expected to rise for many species (Urban 2015). Accurately predicting species’ responses to climate change requires understanding how climate affects survival, growth and reproduction (Paniw *et al*. 2021) – traits that shape population dynamics (Ehrlén & Morris 2015). However, building this understanding is challenging, not least because climate can have complex effects on life-histories. Effects of climate may be seasonal (Paniw *et al*. 2019), or vary between populations due to local adaptation (DeMarche *et al*. 2019) or interactions between climatic drivers and other (a)biotic factors (Urban *et al*. 2016). Within populations, responses to climatic variation may differ between demographic groups (Petry *et al*. 2016). Effectively, how climate affects individual fitness can depend on which population, trait, or life-cycle stage is studied (Paniw *et al*. 2021). A lack of long-term demographic data capturing this complexity (Compagnoni *et al*. 2021; Paniw *et al*. 2021) means that our understanding of how climate affects demography and life-history is incomplete, even in comparatively well-studied taxa. In insects, a general lack of demographic data (Salguero-Gómez *et al*. 2016) complicates efforts to relate demographic or life-history traits to climatic variables. This is certainly true of bumblebees (*Bombus* sp.), key pollinators of crops and wild flowering plants in temperate regions, where almost one in four species on the IUCN Red List is in decline (Cameron & Sadd 2020).

Bumblebee losses are driven by multiple factors (Cameron & Sadd 2020; Wilfert *et al*. 2020), including climate change. Bumblebees are largely cold-adapted (Woodard 2017) and when temperatures exceed species’ thermal limits, bumblebee patch occupancy declines (Soroye *et al*. 2020). Moreover, bumblebees do not appear to move their ranges to mitigate climate change effects (Kerr *et al*. 2015), leading to predictions of pronounced range reductions under global warming scenarios (Martínez-López *et al*. 2021).

Accumulating data on different bee species present a picture of how climatic factors could contribute to extinction risks in our focus group: warming may elevate the metabolic costs of overwintering (CaraDonna *et al*. 2018), disrupt foraging success due to reduced flower availability (Miller-Struttmann *et al*. 2015) and increase asynchrony between bumblebee emergence and flowering times (Pyke *et al*. 2016) for example. But this emerging picture is complicated: there is variation in susceptibility to climate change between closely related bumblebee species (Herbertsson *et al*. 2021; Miller-Struttmann *et al*. 2015). In some cases, moderate warming may have positive impacts on bees across parts of their range (Soroye *et al*. 2020), for example by improving flight performance (Kenna *et al*. 2021). Fine-scale data linking climate and bumblebee health and life-history can add greater insight into the fitness consequences of climatic change and the underlying mechanisms.

Here, we use a long-term dataset (2000 – 2014) to study how climate affects *Bombus terrestris,* social insects with an annual life cycle (**Fig 1**). We focus on queens because this reproductive caste spend much of their lives in a solitary state, unable to benefit from colony level thermoregulation that may buffer thermal variation and thus, may be particularly vulnerable to climate change (Woodard 2017). We test how annual and seasonal variation in precipitation and temperature affect key fitness traits in spring queens collected from two populations in northern Switzerland (**Fig 1 & S1; Table S1**). We study these climatic drivers singly, and explore interactions between them. By studying two populations we gain insight into geographic variation in responses, and by linking queen survival and reproduction with a functional trait (body mass) we explore individual heterogeneity in these responses.

**Fig 1.**
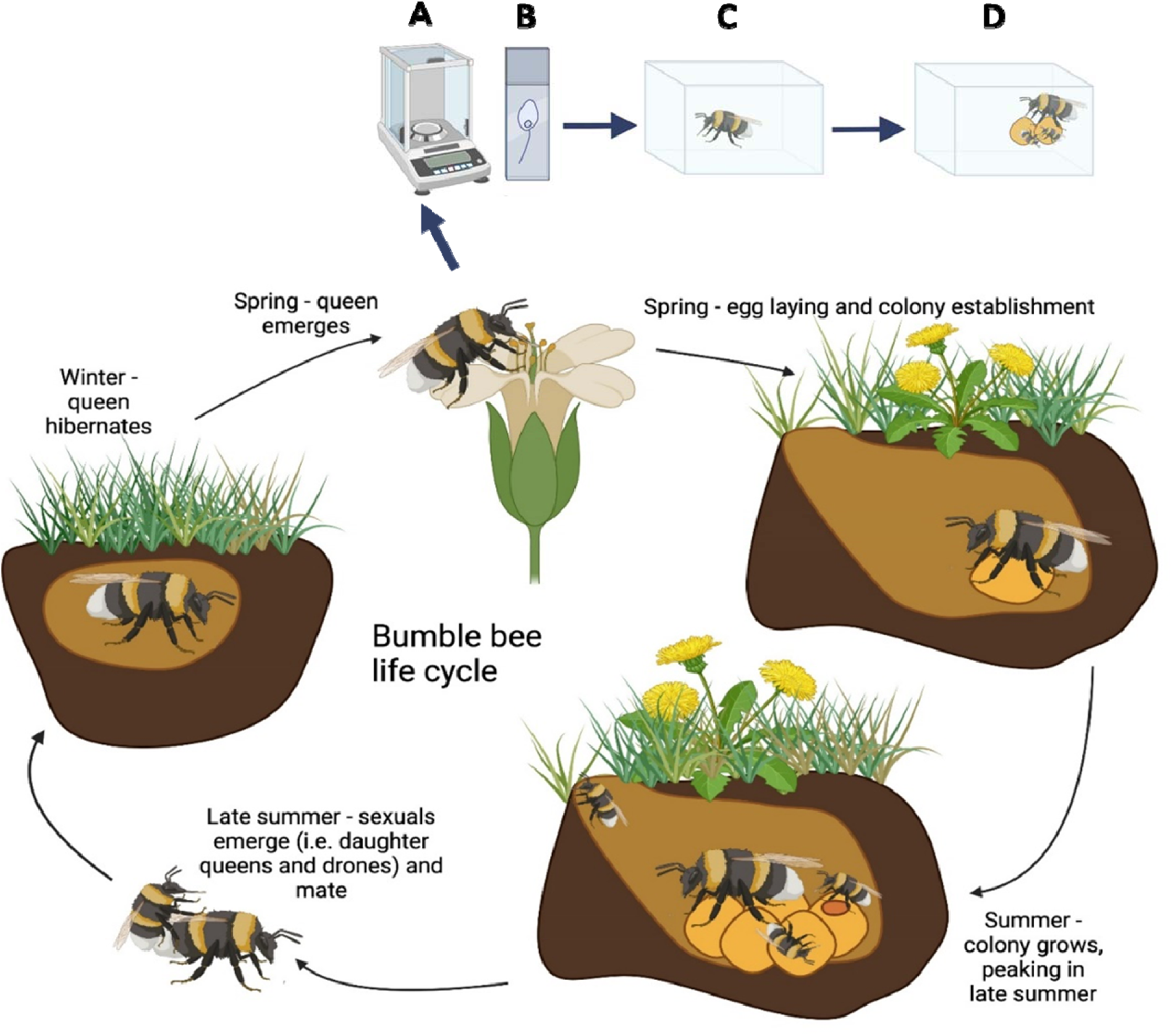
**The annual life cycle of *B. terrestris* and summary of traits measured in the lab (A- D). Life cycle**: Queens emerge from hibernation in the spring, when they consume nectar and pollen and search for a nest site. On finding a site, queens lay eggs that give rise to *a priori* non reproducing daughters (i.e. workers), who begin foraging and help the queen to establish and expand the colony. From this point on, queens remain in the colony and lay eggs that give rise to more workers until summer, when sexuals (i.e. daughter queens and drones) instead of workers are produced. At this time, sexuals leave the colony to mate, whereas the queen and the rest of the colony die. Only the fertilized daughter queens survive and enter diapause, hibernating in the ground until spring, when this cycle restarts with the surviving queens emerging from their hibernation sites. **Data collection**: Queens were collected in spring when they emerged from hibernation. The following traits were then assayed: (**A**) body mass - a marker of overall condition, (**B**) *Crithidia* infection – a gut parasite that reduces *B. terrestris* queen fecundity (Yourth *et al*. 2008), (**C**) survival after emergence from hibernation, and (**D**) success in colony establishment (a key aspect of reproductive success). Mass and parasite infection were assayed immediately after catching queens but survival and colony establishment were measured in queens held under standardized laboratory conditions. At an annual scale, we link these traits to average temperature and precipitation in the year preceding spring queen emergence (i.e. we relate phenotype of a queen collected in year *x* to average climate in year *x* – 1). For greater functional insight, we also use sliding window analyses to relate finer-scale variation in climatic conditions (weekly / monthly data, depending on availability) within a 12- month period preceding the 1^st^ March in the year queens were collected. Figure created in BioRender.

## MATERIALS AND METHODS

### Data collection

4,640 spring queens were caught between 2000 and 2014 from two sites in northern Switzerland – Aesch and Neunforn - for the purpose of establishing experimental lab colonies (**Fig S2; Table S2)**. Every spring, as soon as possible after a warm spell, queens were collected from each site. The few queens caught carrying pollen were released because these were likely to have already established colonies. In the lab, queens were weighed, and their faeces microscopically screened for *Crithidia*. Queens were established in acrylic glass cages (12·5 × 7·5 × 5·5 cm) with *ad libitum* pollen and sugar water. Queens were monitored every 2 to 3 days for egg laying, worker production, survival and cage conditions. Queens that had not reproduced by mid-June (when experimental work typically began) were usually sacrificed by freezing. Not all traits considered here could be assayed in all years or in all individuals – the number of complete cases used to analyse each trait of interest are shown in Table S2. Note that while the vast majority of queens were *B. terrestris,* these data may contain a small number of the sister species *B. lucorum,* which cannot be visually distinguished (Williams *et al*. 2012).

### Weather Stations and Climatic Variables

Weather data were collected from the Swiss Federal Office of Meteorology and Climatology (MeteoSchweiz) portal on the 31^st^ of January 2024. Data for the Aesch study site were collected from the nearby Basel / Binningen Weather station (MeteoSchweiz station abbreviation – BAS).

For the Neunforn site, precipitation data were available from the nearest weather station (Niederneunforn - NIE) but as this station did not collect temperature data, these data were sourced from the nearby Aadorf / Tänikon station (TAE) (**Fig S1, Table S1**). All climatic variables analyzed were standardized relative to historic norms by MeteoSchweiz (**Fig S3-S5**). These standardized variables should be robust to minor variation in absolute temperature or precipitation between each collection site and weather station due to microclimatic or altitude. “Historic norms” are calculated by MeteoSchweiz in 30 year intervals and the operational historic norm used for standardisation was 1991-2020.

Effects of climate were analyzed at two temporal scales. First, spring queen phenotype was related to average measures of temperature and precipitation in the preceding calendar year (i.e., the year the queen was born and raised). Second, to explore seasonal effects, a sliding window approach was used to relate queen traits to weekly (temperature) and monthly (precipitation) climatic variation. Here, we studied a 12-month period before the 1^st^ of March in the year each queen was collected. This period spans the time when the queens in our dataset began emerging from hibernation, and reaches back to when their own mothers exited diapause.

Accordingly, the following climatic variables were used; bold text describes these variables, while abbreviations as used in the MeteoSchweiz website and units are given in parenthesis:

1. annual air temperature 2 m above ground; deviation of the annual mean to the historic norm (tre200yv - °C).
2. **annual precipitation**; **annual total relative to the historic norm** (rre150yv - percentage). Precipitation measures are collected automatically and collection machines are heated, such that snow is included. Annual precipitation and temperature measures were not closely correlated (**Fig S5**).
3. **daily air temperature 2 m above ground; deviation of the daily mean to the historic norm** (tre200dv - °C), aggregated to a weekly level for moving window analyses.
4. monthly precipitation; deviation of the monthly total from the historic norm (rre150mv - percentage). Monthly precipitation was the highest resolution available.

### Statistical Analyses

Analyses were conducted in R version 4.0.5 (R Core Development Team 2020) using the following packages: *dplyr* (Wickham *et al*. 2021) for data wrangling, *lme4* (Bates *et al*. 2015), *car* (Fox & Weisberg 2019), *DHARMa* (Hartig 2021), survival (Therneau 2021), *climwin* (Bailey & van de Pol 2016) and *piecewiseSEM* (Lefcheck 2016) for analyses and model checking and *gridExtr*a (Auguie & Antonov 2017) and *ggplot2* (Wickham 2016) for plotting.

Data were analyzed in three steps:

**1.** Individual mixed models related each trait to annual climatic measures in the preceding calendar year.
**2.** Piecewise structural equation modelling was used to test for complex relationships between multiple interrelated variables and annual climatic variation.
**3.** A sliding window approach related weekly (temperature) or monthly (precipitation) data to spring queen phenotype (March – March).

This approach balances the risks of multiple testing (i.e. studying traits collected in the same animals in separate models) against the loss of power due to a reduction in sample size if only complete observations are analyzed in a single piecewise SEM.

#### Step 1) Spring Queens and Annual Climatic Variation

Individual mixed models were constructed for each trait. Study site (factor, two levels), collection day of the year (continuous) and queen body mass (continuous) were always included as covariates (excluding body mass when that was the response trait). Continuous variables were scaled to have a mean of zero and a standard deviation of one to facilitate model convergence. Year (factor) was modelled as a random intercept. For each trait, linear effects of temperature and precipitation were included in models, as was the interaction between them. For *Crithidia* and reproduction binomial generalized linear mixed models (Bates *et al*. 2015) were used with the *bobyqa* optimizer to facilitate convergence. For body mass, a linear mixed model was used with a Gaussian error structure. *DHARMa* was used to check model fit. For binomial models, fit was assessed using a “grouping” method, which turns 0/1 Bernoulli data into a k/n response.

Survival was analyzed using mixed cox-proportional hazards models and deviation from the proportionality assumption was tested using the function *cox.zph* (for detail see Text S1). The Anova function from the *car* package was used to test significance in the full models.

#### Step 2) Piecewise SEM

Piecewise structural equation modelling using the function *piecewiseSEM* (Lefcheck 2016) was used to evaluate causal relationships between climatic variables, functional traits and reproduction. Survival was not included because survival models cannot be incorporated into this modelling framework. The starting *piecewiseSEM* model is shown in **Fig S6.** As for the annual models described above, continuous explanatory variables in this starting model were scaled (bar body mass, which was also a response variable in this model), and year was included as a random factor. After creating this starting model, we used d-separation tests to determine if any paths that were not specified in the starting model were significant and Fishers c-tests to evaluate model fit. We report conditional (*R*^2^) coefficients for each of the models incorporated into the overall *S*EM, because this measure considers variance associated with fixed and random effects.

#### Step 3) Sliding window analyses

Sliding window analyses were performed using the *climwin* R package to identify periods of time when effects of climate on each trait were most pronounced. Our baseline model set the 1^st^ of March in the year each queen was collected as a reference date, and related climate in year preceding this reference date to traits of interest. Our baseline model included collection day of the year (continuous, scaled), and body mass (continuous, scaled; except where body mass was the response trait). Year (factor) could only be included as a random effect in analyses of continuous traits (i.e. body mass) and in other cases, was included as a main effect. We tested for linear and quadratic climatic effects on each trait and site was accounted for using the “spatial” term. Queen phenotype data from 2000 was excluded for temperature, due to the appropriate climatic variable not being available for both sites in 1999. We included body mass as a linear term for consistency and model output did not change qualitatively if mass was also included as a quadratic term for reproduction (see Results). Randomization analyses were used to quantify the likelihood of obtaining the level of model support observed for the best model by chance alone. We report *P* values for these analyses calculated using 500 randomizations and degrees of freedom equivalent to the number of years multiplied by the number of study sites.

Where more than one model had a similar probability of being the best model in the model set, we report the best model and used the confidence set - a subset of models with 95 % confidence of including the true best model – to calculate median windows. We used model averaging to calculate the average relationship between climate and traits in the model set, weighted by the probability that each model in the set is the best model. For survival, the top performing models in the confidence set suggested multiple climatic peaks. In this case, we split the year up into three ranges, each encompassing a candidate peak, and repeated the analyses described above for each range.

## RESULTS

### Body Mass

Collection day did not impact body mass (Χ^2^ = 2.26, df = 1, *P* = 0.133) but site did: queens collected from Neunforn were ca. 4% heavier than queens collected from Aesch (Χ^2^ = 62.78, df = 1, *P* < 0.001). Temperature and precipitation interacted to affect body mass (Χ^2^ = 7.26, df = 1, *P* = 0.007) (**Fig 2**): when temperatures were relatively low, precipitation had a positive effect on body mass but effects of precipitation became increasingly negative with increasing temperature. As a result, warm, wet years correlate with lower body mass. The independent main negative effect of temperature (correlating with lower body mass; Χ^2^ = 3.49, df = 1, *P* = 0.062) and positive main effect of precipitation (correlating with larger body mass; Χ^2^ = 2.84, df = 1, *P* = 0.092) were approaching significance. Outputs of *piecewiseSEM* matched these annual modelling results (**Fig 3**). While global fit statistics suggested that the piecewise SEM fit the data, this model explained relatively little variation in body mass (conditional R squared – 0.06). At a finer temporal resolution, moving window analyses highlighted that elevated precipitation between May and November, and a warm period in November reduced body mass (**Fig 4; Fig S8; Table S3-S4**).

**Fig 2.**
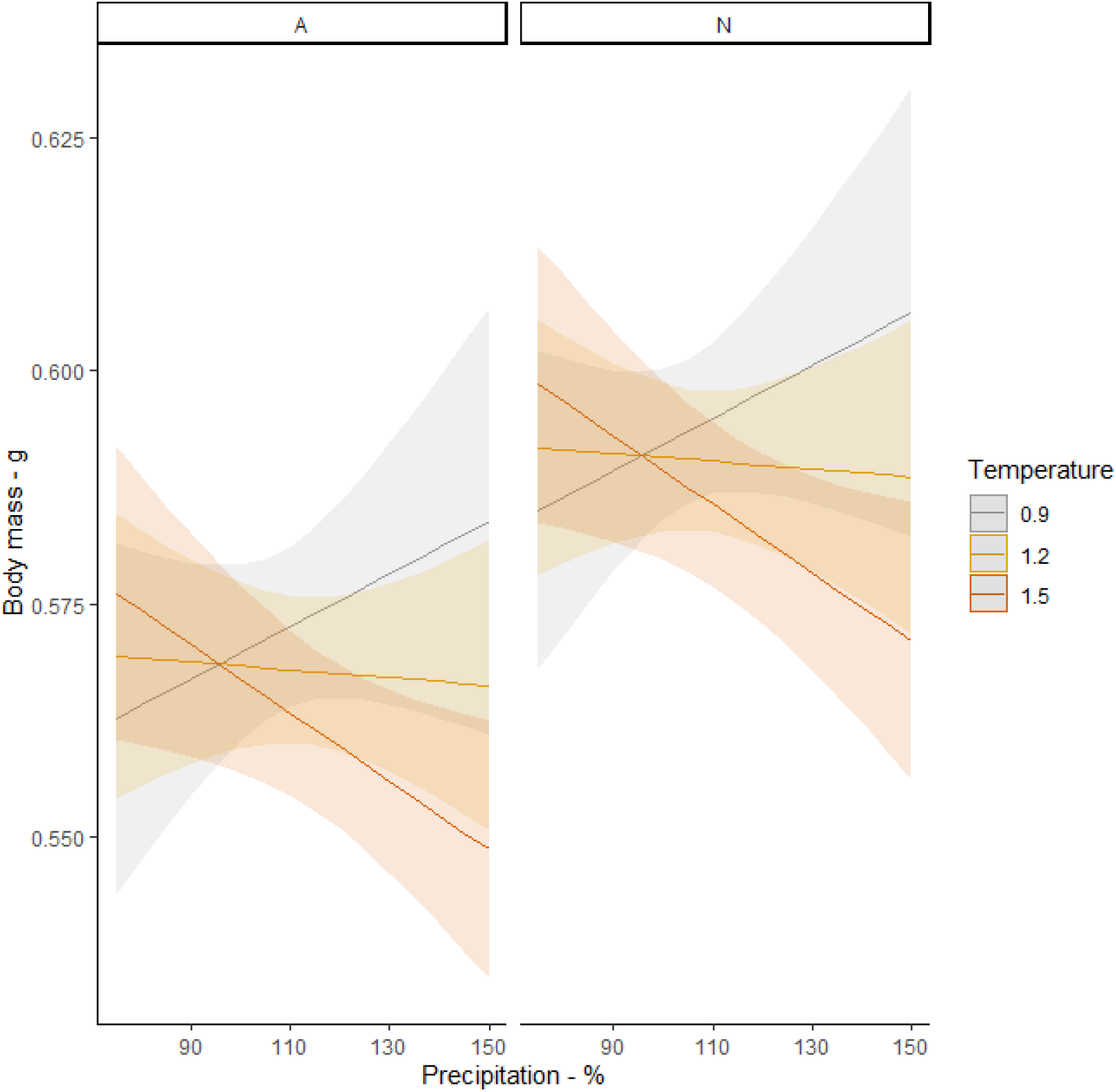
The predicted interaction effect between temperature and rainfall in the preceding year on body mass in Aesch (A) and Neunforn (N). Shaded regions show confidence intervals, and lines show predictions from the model estimated. To plot the interaction, the relationship between precipitation and body mass is shown at the lower, middle and upper terciles of temperature variation – 0.9°C (grey), 1.2 (yellow) and 1.5 (red) respectively. These temperatures represent the terciles of the standardised temperature variable - lower, middle and upper respectively. Precipitation is given as percentage deviation relative to the historic norm.

**Fig 3.**
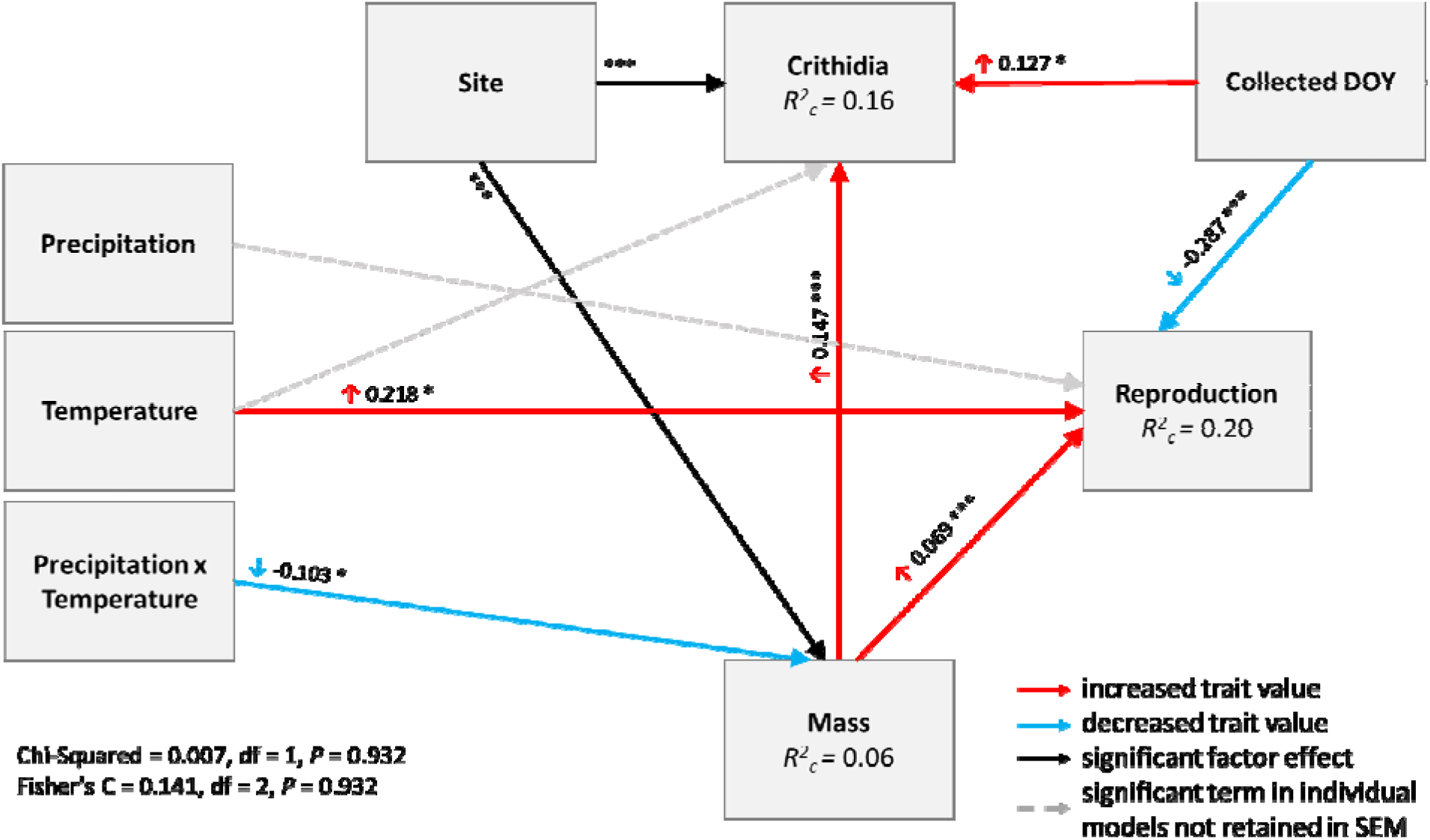
Structural equation modelling. Arrows show significant effects and the direction of these effects, with red arrows showing positive and blue arrows showing negative relationships. Black arrows are used for site effects. Grey dashed arrows highlight differences between the outcomes of this model and the annual analyses i.e. terms that were significant in the annual models but not in the piecewise SEM approach. Standardized path coefficients are reported next to each of the arrows, with colour coded arrows shown to make it easier to see the direction of effects. R^2^ values (conditional R^2^c) are reported within the boxes for each of the continuous response variables. Note that only significant relationships are shown here for clarity, where * = P ≤ 0.05; ** = P ≤ .01; *** = P ≤ 0.001.

**Fig 4.**
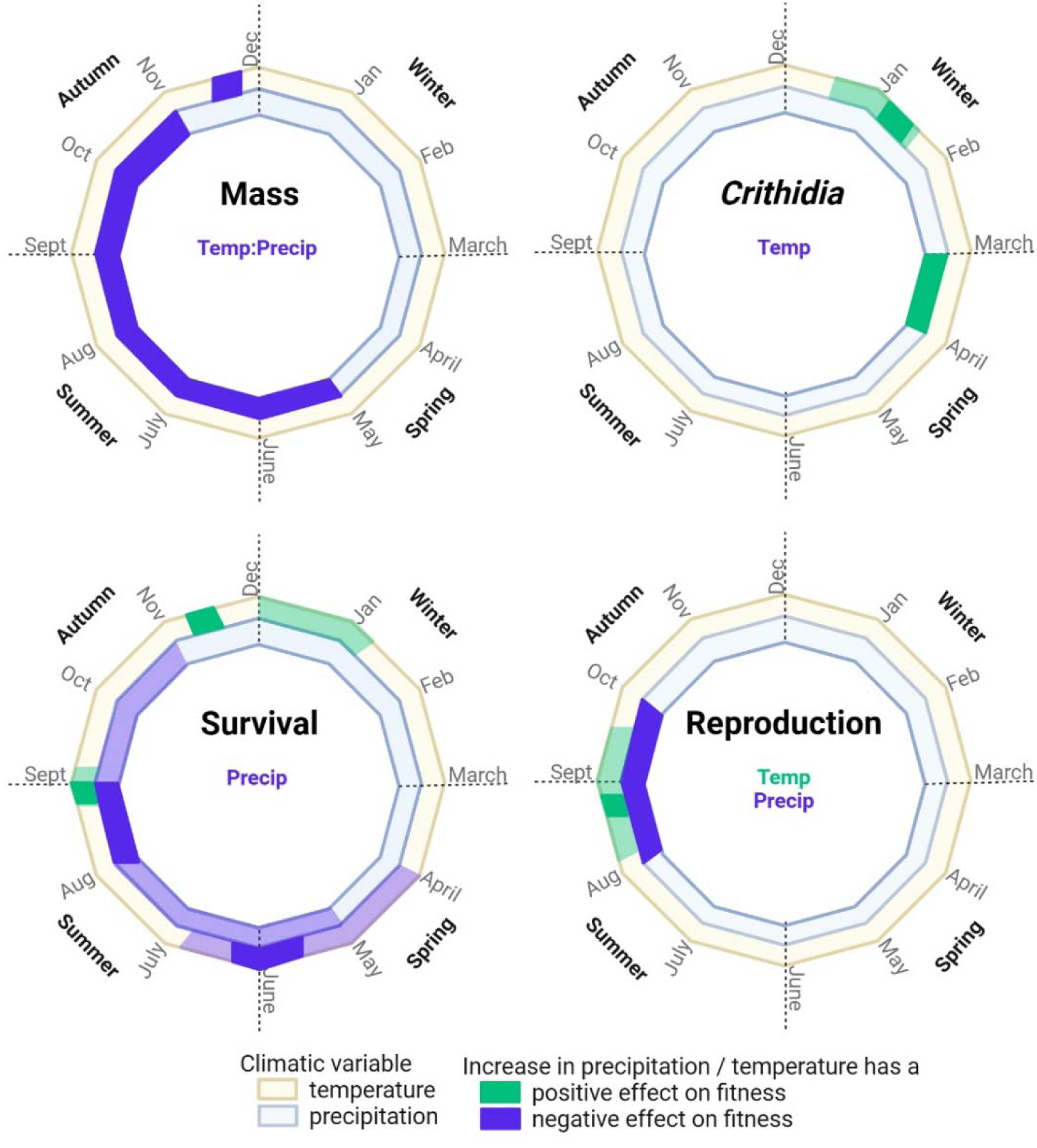
Summary of sliding window analyses of body mass, *Crithidia* infection, survival and reproduction. Each ring is divided into monthly steps to represent the entire year covered in the sliding window analyses. The outer yellow coloured ring displays results for temperature (weekly steps) and the inner blue ring shows results for precipitation (monthly steps). Month labels show the 1^st^ of each month, and the year divided using dashed lines into seasons. The writing inside each circle summarises the results for the annual analyses – only significant climatic effects are shown. Colour coding shows the effect of a rise in each climatic driver on fitness, where green means a positive change in fitness and purple a negative change. Light colours indicate windows identified in the confidence set, and dark colour windows identified in the single best model. Note that the direction of the effect on fitness may oppose the direction on the trait i.e. survival analyses models hazard (i.e. the risk of dying), therefore an increase in hazard is detrimental to fitness. These schematics should be interpreted alongside precise estimates of window duration in **Table S3-S4**.

### Crithidia

7% of queens from Neunforn were infected with *Crithidia* but 13% of Aesch queens were infected – a significant difference (Χ^2^ = 43.10 df = 1, *P* = <0.001). The prevalence of infection was higher in heavier queens (Χ_2_ = 18.01, df = 1, *P* = <0.001), and in queens collected later in the year (Χ^2^ = 5.52, df = 1, *P* = 0.019). Precipitation did not affect *Crithidia* prevalence singly (Χ^2^ = 2.17, df = 1, *P* = 0.141) or via its interaction with temperature (Χ^2^ = 0.12, df = 1, *P* = 0.733). However, queens collected after an above-average warm year were more likely to be infected (Χ^2^ = 5.50, df = 1, *P* = 0.019) (**Fig S7**). While effects of site, body mass and collection date were conserved between analytical approaches, the piecewise SEM analysis suggested that climate did not directly affect *Crithidia* incidence (**Fig 3**). Instead, effects of climate on *Crithidia* were mediated via changes in body mass.

Moving window analyses suggested significant impacts of temperature and precipitation on *Crithidia* incidence (**Fig 4, S9, Table S3-S4**). The best fitting model suggested that the likelihood of *Crithidia* infection was reduced by warm temperatures in mid-January. Similarly, a reduction in *Crithidia* prevalence was suggested by averaging models in the confidence set, in particular, by warm temperatures from late in December to mid-January. Note that these analyses suggest that warm winters reduce infection but overall warm years (see above) increase infection.

Analyses of precipitation data suggested that wet conditions in March in the year preceding spring queen emergence (when the mothers of focal queens were emerging and establishing their colonies) reduced *Crithidia* incidence.

### Reproduction

The fit of the reproduction model was initially poor, and improved by including a quadratic effect of mass (Χ^2^ = 9.25, df = 1, *P* = 0.002). This means that while heavier queens were more likely to establish colonies (i.e. positive linear effect of mass, Χ^2^ = 11.70, df = 1, *P* = 0.001), the benefits of being large had diminishing returns. Colony establishment was independent of site (Χ^2^ = 0.02, df = 1, *P* = 0.892) but greater in queens collected earlier in the season (Χ^2^ = 25.98, df = 1, *P* = <0.001). Temperature (Χ^2^ = 6.65, df = 1, *P* = 0.010) and precipitation in the preceding year (Χ^2^ = 5.41, df = 1, *P* = 0.020) affected colony establishment; warm preceding years increased the likelihood of reproduction, while wet years reduced it (**Fig 5**). Interactions between climatic variables did not significantly affect reproduction (Χ^2^ = 0.50, df = 1, *P* = 0.482). Piecewise SEM analyses largely agreed with these results, but not entirely. In this global model, there was no significant direct effect of rainfall on reproduction. Instead, rainfall affected body mass via an interaction with temperature, and this in turn affected reproduction (**Fig 3**).

**Fig 5.**
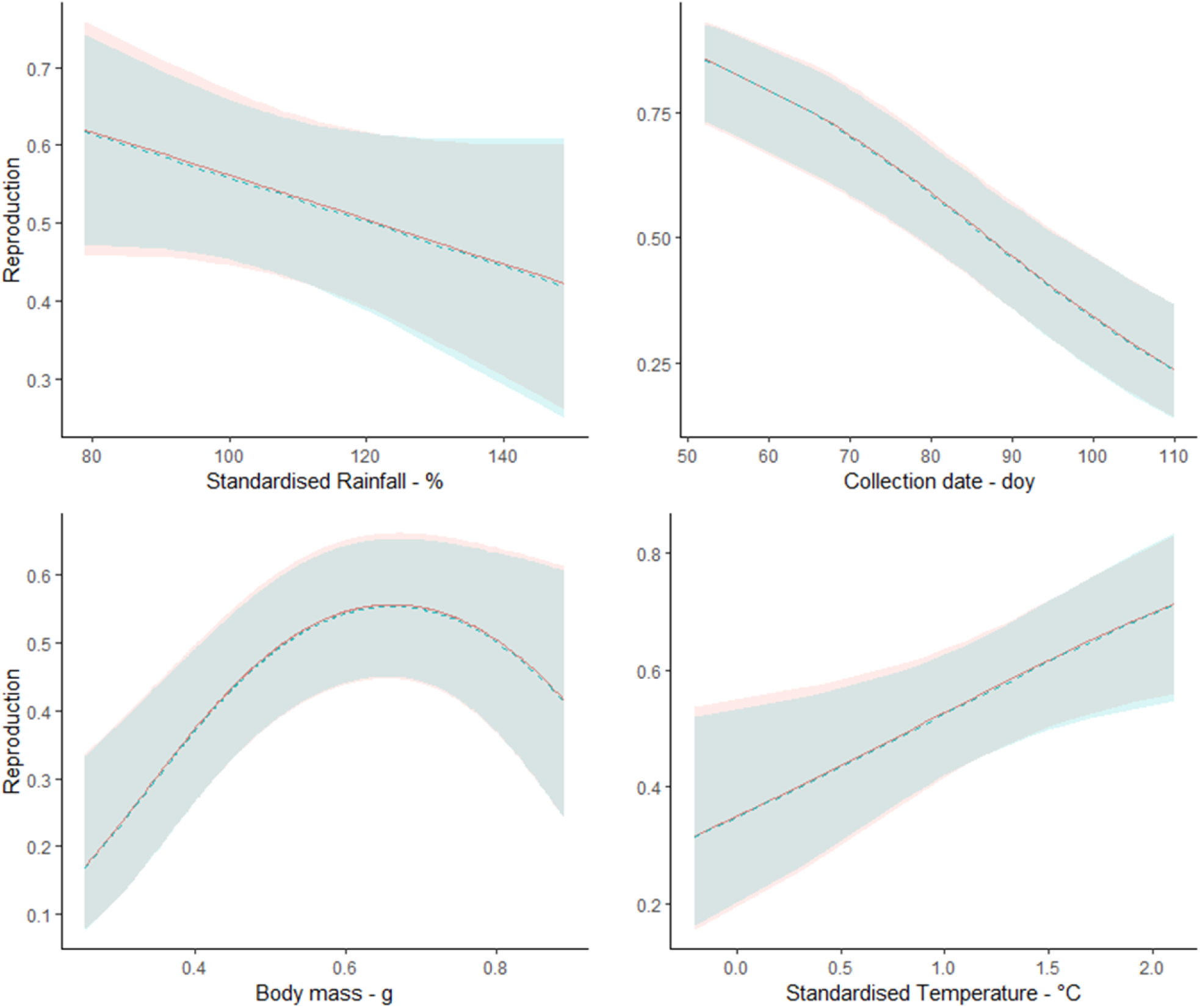
The predicted relationship between the likelihood of successful reproduction (i.e. colony establishment) and queen body mass (g), the day of the year (DOY) when queens were collected, standardised rainfall (% – linear) and standardised temperature (C - linear). Shaded regions show confidence intervals, and lines show predictions from the model estimated using the *ggpredict* function in the *ggeffects* package. Red = site Aesch, Blue = site Neunforn.

The best model from moving window analyses suggested that warm Augusts increased the likelihood of reproduction, although the broader confidence set suggested that a wider period - spanning August and September - was particularly important. Pronounced precipitation in August and September appeared to reduce subsequent reproduction (**Fig 4, Fig S10, Table S3- S4**).

### Survival

The risk of dying (i.e. the hazard ratio) was lower in heavier queens (coefficient ± standard error: -1.54 ± 0.57, Anova: Χ^2^ = 7.35, df = 1, *P* = 0.007). Queens collected later in the season (coefficient ± standard error: 0.30 ± 0.09, Anova: Χ^2^ = 11.63, df = 1, *P* = <0.001) and after wet (preceding) years (coefficient ± standard error: 0.18 ± 0.12, Anova: Χ^2^ = 9.99, df = 1, *P* = 0.002) had a greater risk of dying. Neither site (Χ^2^ = 2.24, df = 1, *P* = 0.134) nor temperature (Χ^2^ = 3.04, df = 1, *P* = 0.081) significantly affected this risk. That said, temperature effects bordered on significance, suggesting a tendency towards higher death hazard following warmer conditions (coefficient ± standard error: 0.20 ± 0.11). Finally, the interaction between climatic variables did not affect hazard (Χ_2_= 0.61, df = 1, *P* = 0.434)

The single best model produced by moving window analyses suggested that high levels of precipitation in August relative to historic norms elevated the risk of dying. The broader confidence set suggested that precipitation between May and October was particularly detrimental (**Fig 4, Fig S11, Table S3-S4**). Seasonal effects of temperature were complex. The first model relating temperature across the entire year to mortality risk identified three influential climatic windows (**Fig S11**). Thus, further models were run for periods corresponding to each peak (1 – 19 weeks prior to the 1^st^ of March; 20 – 29 weeks and 30 – 52 weeks). The first period focussed on temperatures between mid-October and March 1^st^ in the year that spring queens were collected. The best model for this period suggested that a warm spell early in November decreased the risk of dying. However, analyses of the subset of models from the confidence set identified a period of time that did not overlap with the output of the single best model, which reduces our confidence in this model output. Despite this, randomization analyses suggested that this “best model” was unlikely to arise by chance alone (**Table S4**).

The single best model for the second period (i.e. a second step back in time, 20 – 29 weeks prior to the 1^st^ of March), suggested that warm periods spanning late August to early September reduced the likelihood of spring queens dying prior to reproducing the following spring. The broader model set encompassed this single best model but suggested that the influential window of time was broader, reaching into September (**Table S4**).

Analyses of the third time period (30 – 52 weeks prior to the 1^st^ of March), suggested that high temperatures in mid-May to early June elevated mortality risk. The confidence set suggested that high temperatures from April until mid-June significantly elevated mortality risk (**Table S4**).

## DISCUSSION

Each spring, *B. terrestris* queens emerge from hibernation and establish their own colonies, producing workers and then daughter queens, who forage, mate, hibernate and then (perhaps) emerge to begin this cycle again. Using long-term data, we link variation in climate across this annual cycle to multiple fitness traits in queens that survive diapause to emerge in spring. In this cohort of survivors, average climatic conditions across the preceding year affected almost all fitness traits. Queens emerging after wet years displayed consistent reductions in fitness but in contrast, emerging after warm years had positive and negative fitness impacts via effects on different traits. Beyond these annual patterns, there was pronounced seasonality in the effects of climate, revealing when in their lifecycle queens are most vulnerable to climatic variation – in particular, when queens are produced late in summer, forage, mate and enter hibernation. In line with established knowledge on this system, these results suggest that climatic conditions that reduce resource acquisition in summer, or exacerbate resources depletion overwinter, are vital determinants of spring queen fitness.

At an annual scale, elevated precipitation was largely detrimental to spring queen fitness. Queens emerging after a wet year were less likely to survive and reproduce, and warm-wet years reduced spring queen body mass. Warming also had negative fitness effects, because queens emerging after warm-wet years were lighter, and more likely to be infected with *Crithidia*. However, warming in the preceding year had a non-significant impact on survival and even improved the likelihood that spring queens established their own colonies. This suggests that warming might have positive impacts on the fitness of spring queens. This is possible because moderate warming can improve traits such as bumblebee flight (Kenna *et al*. 2021).

However, the apparent benefits of emerging after a warm year are not as positive as they first appear. Structural equation modelling showed that while warming increased colony establishment, warm-wet years reduced body mass and lighter bees were less likely to establish colonies. That is, warming had a positive direct impact on the colony establishment and a negative indirect effect. The direct negative effect of precipitation on reproduction was not recovered in this structural equation model, suggesting that precipitation effects on colony establishment may also be mediated via body mass. Similarly, in this global model, climate did not directly affect parasite prevalence, but the interaction between temperature and precipitation reduced body mass, and lighter bees were less likely to host *Crithidia*. Evidently, some fitness effects of climate on spring queens are mediated via effects on body mass.

That mass mediates effects of climate on queen fitness is not surprising because body mass affects many aspects of bumblebee health and performance. Heavier *B. terrestris* workers have greater nectar foraging rates (Spaethe & Weidenmüller 2002), colonies with larger nurses produce more workers (Cnaani & Hefetz 1994) and queens with a wet weight below < 0.6 g are unlikely to survive diapause (Beekman *et al*. 1998). This latter benefit is important in this dataset, where mass was measured only in queens that successfully overwintered and lighter queens may not have survived winter to feature in the dataset at all. This selective disappearance of lighter queens may mean that the importance of body mass on bumblebee fitness is underestimated in these analyses – the fitness of light queens is null if they die overwinter. If the severity of selective disappearance overwinter depends on climate, these data may also underestimate how much climate affects body mass. Crucially, the significance of body mass as a direct or indirect mediator of climatic condition observed here, can justify this measure as a simple, practical tool for studying the effects of climate change.

Climate effects on the mass of queens emerging from hibernation may reflect that climatic factors affect resource acquisition during queen development, or resource loss overwinter. Body mass certainly speaks to resource availability during development, as young queens gain body mass rapidly in the first 2-3 days of their adult life, with mass acquisition subsequently slowing down (Woodard *et al*. 2019). Queens undergoing diapause are exposed to low temperatures and food limitation and therefore display pronounced reductions in their body mass and lipid reserves (Ghosh *et al*. 2017). These reserves affect survival during diapause, and reserves remaining after diapause determine how readily individuals emerging from diapause can invest in fitness traits, such as reproduction (Hahn & Denlinger 2007). Sliding window analyses suggests that climate affects spring queens via both routes i.e., by modulating mass gain in summer and mass loss overwinter.

A clear signal that climate exacerbates mass loss overwinter is that warm winters reduced spring queen body mass. This mirrors data from *Osmia ribifloris*, an early-season solitary bee, where experimental warming of field populations was associated with reductions in body mass (CaraDonna *et al*. 2018). Similarly, *B. lucorum* queens emerging after induced diapause in warm conditions in the lab, tended to have less body fat then queens that overwintered at lower temperatures (Vesterlund *et al*. 2014). The most likely explanation for these results is that warm winters encourage queens to emerge from their quiescent state and underground habitats, elevating metabolic costs at a time when forage is rare, therefore reducing fat stores and mass at emergence. Thus, some of the detrimental phenotypic effects of warming observed in these data seem likely to be due to elevated metabolic costs of overwintering.

Warm winters also appear to reduce mortality risk and *Crithidia* incidence in spring. If elevated metabolic costs of surviving warm winters reduce body mass – a key fitness trait - how can they also reduce disease incidence and mortality? The answer to this apparent contradiction may be selective disappearance. *Crithidia* exacerbates weight loss in hibernating queens (Brown *et al*. 2003), and because body mass is an important predictor of overwinter survival (Beekman *et al*. 1998), small and infected queens may not have been able to tolerate this high metabolic cost. These queens may have died before emerging in spring to feature in our dataset, meaning that any surviving infected queens tend to be bigger on average. Similarly, if warm winters elevate overwinter mortality and mortality is condition dependent, then the queens in this dataset may be likely higher quality than average and perhaps, more likely to survive subsequently. Data testing how climate affects the condition and health of queens entering hibernation and those emerging from it would offer some insight into whether selective disappearance and condition dependent mortality explain these apparently counterintuitive results. At least for infections by another parasite, the microsporidian *Nosema bombi,* the (mitochondrial) haplotype of spring queens may additionally affect observed prevalence with respect to climate variables (Manlik *et al*. 2023). This may add further complexity to the study of disease risk under climate change.

Another sign that climate may exacerbate overwintering costs is that warm and dry conditions around August and September improve survival and reproduction. In mountainous populations in south-eastern Germany queens are rarely observed foraging (and thus may have begun hibernating) after July (Sponsler *et al*. 2022). This means that at some point in this critical window of time, many queens in our Swiss population may have entered hibernation. Warming late in summer may also delay the onset of diapause and thus, could reduce its metabolic costs. This rationale may also explain why queens collected later in the spring sampling period were less likely to survive and reproduce. While not explicitly a measure of climate, later collection dates may reflect that queens have experienced cold springs, or springs characterised by oscillating temperatures, that extend hibernation and so further elevate the metabolic costs of surviving winter and deplete reserves needed to successfully establish colonies in spring.

Although of course, there may be costs to emerging from diapause too early, if forage is not yet available.

There are also signs that climate reduces mass acquisition during queen development. Wet conditions in spring, summer and autumn reduced body mass and spring queen survival for example. The negative effect of wet springs and summers may reflect reduced resource acquisition, because rainfall reduces foraging activity and high humidity limits pollen retrieval (Karbassioon *et al*. 2023; Reeves *et al*. 2023). Similarly, the positive effects of warm and dry conditions around August and September on survival and reproduction could be due to effects of climate on resource acquisition and / or expenditure rather than on delayed hibernation.

While mountainous populations may be beginning to hibernate during this time, in populations in southern Germany and in the UK, new queens and drones are often on the wing (Frank 1941; Haas 1946) (pers. comm. Richard Comont (UK BeeWalk) and Hannah Burger (Germany, BienABest)), foraging to amass resources for hibernation. Warm, dry conditions during this time may improve forage availability, at a time when there may be a nectar deficit for bumblebees (Timberlake *et al*. 2019). In support of this idea, late-summer nectar availability predicts colony density in the following year (Timberlake *et al*. 2021). Warm, dry conditions late in summer could also improve foraging success because rainfall reduces bumblebee foraging activity (Reeves *et al*. 2023) and at low temperatures in a range that may be increasingly common in early autumn (i.e. 12°C), only large bees can fly (Kenna *et al*. 2021). By affecting forage availability or foraging efficiency, warm-dry periods late in summer could elevate queen resources before they enter hibernation when they have to survive overwinter largely on their accumulated resources.

Warming and increased rainfall are already having costly impacts on spring queen fitness even in our study populations, which are not near their upper thermal limits, with body mass a likely mediator of effects. While there are many candidate mechanisms that could explain the patterns identified here, we suggest that climatic factors that make it harder for queens to acquire resources in summer, and exacerbate resource loss overwinter, are particularly detrimental. If correct, this means that climate change and forage quality are likely interacting to affect bumblebee queen fitness. This is not surprising given that nutrient intake is a central determinant of female fitness in insect species (Bunning *et al*. 2016; Carey *et al*. 2022; Rapkin *et al*. 2018), including bumblebees (Stabler *et al*. 2015; Vaudo *et al*. 2016). If forage influences effects of climatic change, then climate effects may be acting in synergy with land-use intensification, which also affects body mass and resource acquisition in bumblebees (Straub 2023). While warmer winters and interrupted springs can only be addressed by global efforts to counter climate change, providing more floral resources during crucial windows of colony development may mitigate effects of climate change and land use intensification, including in other pollinator species (Timberlake *et al*. 2019).

## Data Accessibility Statement

**Climate Data** - Data are subject to third party restrictions but are available on request from MeteoSchweiz. **Phenotypic Data** – Data will be uploaded to an open access repository at manuscript acceptance. For review, all data and accompanying R code have been uploaded with the manuscript. This is because we are not allowed to submit climate data to an open-access repository (restriction imposed by MeteoSchweiz) but want Reviewers to have access to the full data bundle.

### Acknowledgements

We would like to thank everyone involved in data-collection, in particular Rahel Salathé, Christine Reber Funk, Roland Loosli, Chrissy Gerloff, Elmar Benelli, Yannick Moret, Ben Sadd, Oliver Otti, Mark Brown, Pius Korner, Boris Baer, Hauke Koch, Yuko Ulrich, Seth Barribeau, Chris Yourth, Martina Tognazzo, Gabriel Cisarovsky, Dani Heinzmann. We would like to thank owners of the sites where bees were collected, MeteoSchweiz for climatic data, Hannah Burger and Richard Comont for sharing information on queen bee flight periods. Finally, thanks to Martijn van de Pol and Liam D Bailey (climwin) and Jonathan S. Lefcheck (piecewiseSEM) for advice using their statistical packages and Erik Postma and the Wilfert lab group for helpful discussion. CRA is funded by DFG grant - AR 1369/2-1. PSH was funded by Swiss National Science Foundation: grants nos. 31-49040.96, 31-66733.01, and 3100A-116057; intra-mural funding: CCES ERU 0-21220-07.

## Author Contributions

CRA performed analysis and wrote the first draft. LW and RSH collected data. PSH, LW and RSH edited the manuscript. All authors helped conceive manuscript.

