## Supplementary material for "WARMING AND INCREASED RAINFALL REDUCE BUMBLEBEE QUEEN FITNESS": Online Supplement

**CONTENTS**

| PART 1 – STUDY SITE LOCATION | Page 2 |
| --- | --- |
| PART 2 – COLLECTION DATES AND SAMPLE SIZES | Page 4 |
| PART 3 – SITE CLIMATIC INFORMATION | Page 6 |
| PART 4 – EXTRA DETAIL STATISTICS | Page 10 |
| PART 5 – EXTRA INFORMATION ANNUAL RESULTS | Page 13 |
| PART 6 – EXTRA INFORMATION SLIDING WINDOWS | Page 14 |

**PART 1 – STUDY SITE LOCATION**

**Fig S1. Map of northern Switzerland showing the location of study sites and associated weather stations.** Closed circles show the study sites (red – Aesch, green – Neunforn), while open circles show the associated weather stations. For Neunforn, weather data originated from two sites: precipitation data were provided by the nearby Niederneunforn station, while temperature data originated from the Aadorf / Tänikon weather station, which is south-east of the study site.

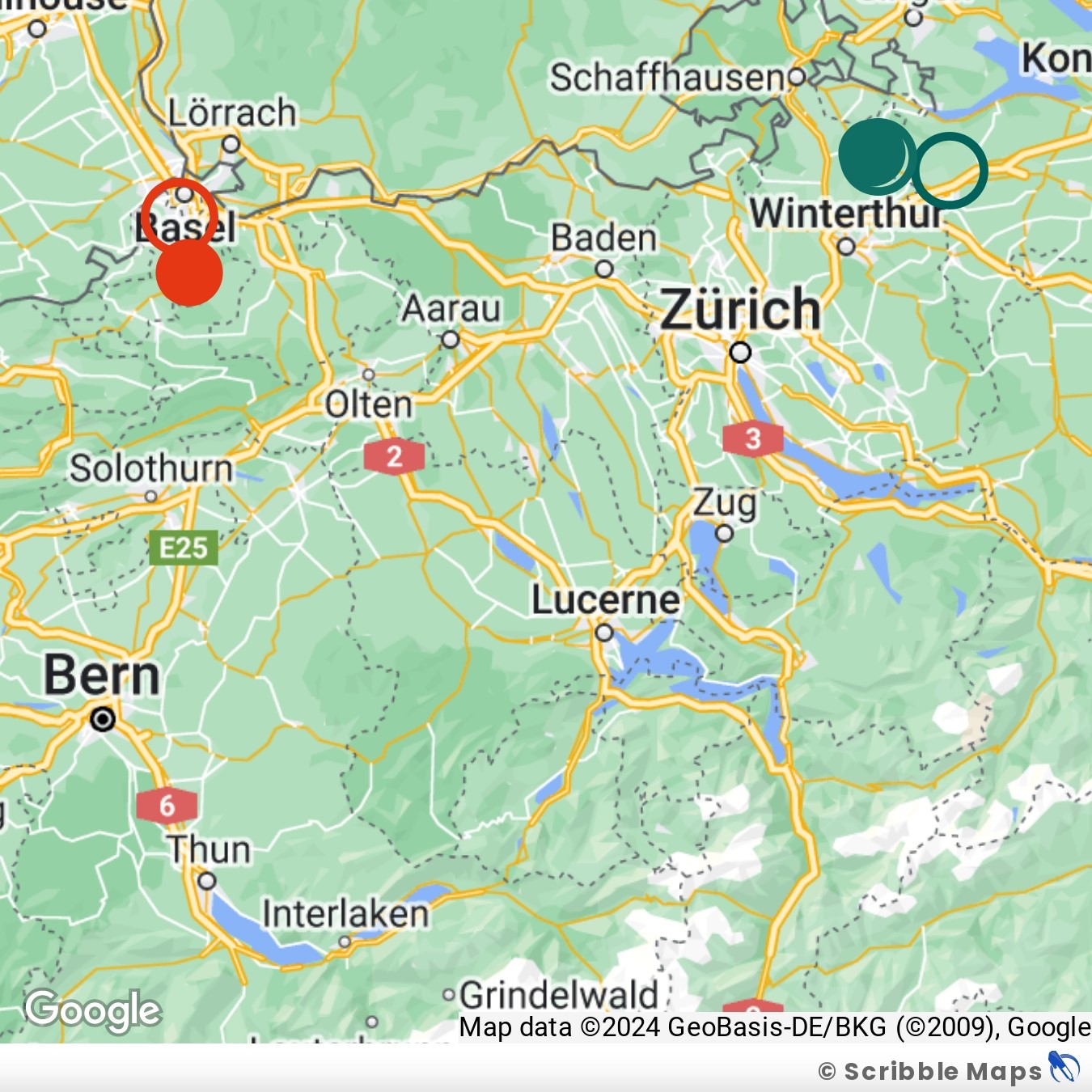

**Table S1. Location information for each study site and weather station that provide climatic data.** Geographic information for each weather station are exact and provided by MeteoSchweiz, while geographic data correspond to the approximate location where bees were collected.

| Location | Description | m | Longitude / Latitude |
| --- | --- | --- | --- |
| Aesch | Study Site | 309 | 7° 34′ / 47° 28′ |
| Basel / Binningen | Weather station | 316 | 7°35' / 47°32' |
| Neunforn | Study Site | 460 | 8° 46′ / 47° 36′ |
| Aadorf / Tänikon | Weather station | 539 | 8°54' / 47°29' |
| Niederneunforn | Weather station | 440 | 8°47' / 47°36' |

**PART 2 – COLLECTION DATES AND SAMPLE SIZES**

**Fig S2. Variation in queen collection dates in each year of sampling.** The date of the year (1 = January first) when each queen in the data-set was collected, presented by year. Colours group points by sampling year. The 1^st^ of March is the reference date for sliding window analyses, and is shown as an inverted filled triangle on the x-axis.

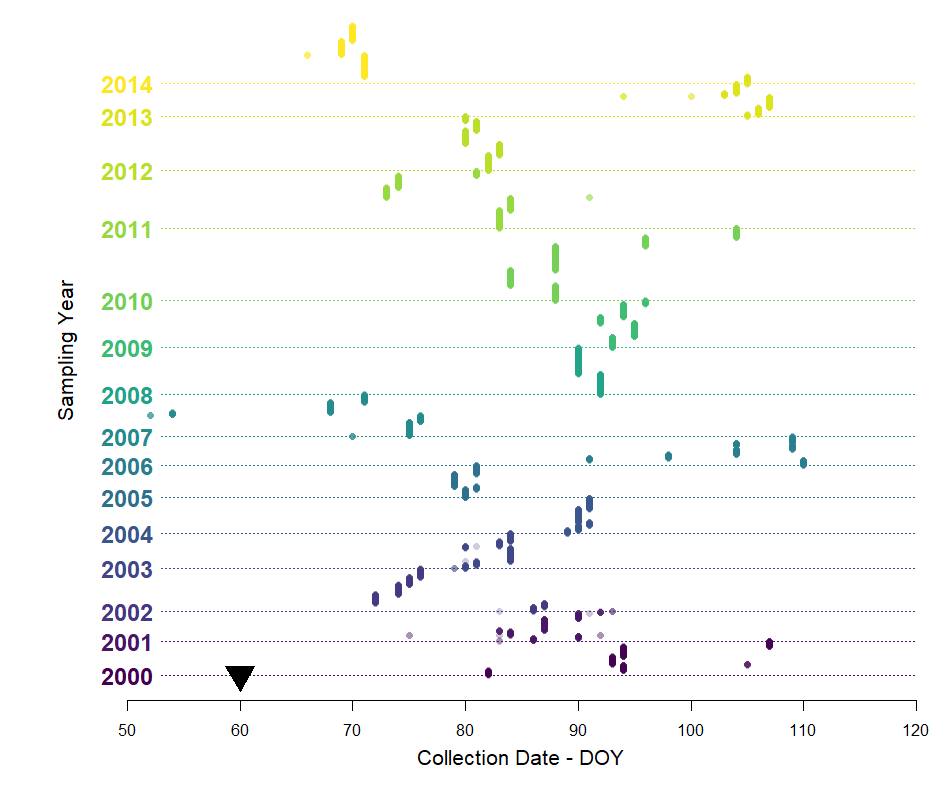

**Table S2. Trait Specific Sample Sizes**. The total column refers to the number of queens sampled. However, because not all traits were measured in all animals, and in all years, sample sizes differed between traits of interest. The numbers below reflect the number of “complete cases” that were used in the model for each trait i.e. observations where data were available for the response variable of interest and relevant covariates (mass, collection date). For example, the *Crithidia* column, shows the number of queens collected that were assayed for Crithidia, and had information on collection date and body mass. Path stands for pathway analyses. In 2012 and 2013 mass was not recorded, because this is a vital co-variate that includes expression of all other traits assayed, queens lacking mass information were not included in analyses.

| Year | Total | Crithidia | Mass | Survival | Reproduction | Path |
| --- | --- | --- | --- | --- | --- | --- |
| 2000 | 243 | 243 | 243 | 130 | 242 | 242 |
| 2001 | 212 | 203 | 209 | 66 | 167 | 162 |
| 2002 | 304 | 248 | 303 | 133 | 139 | 116 |
| 2003 | 254 | 244 | 247 | 230 | 235 | 232 |
| 2004 | 253 | 250 | 253 | 249 | 253 | 250 |
| 2005 | 233 | 214 | 232 | 180 | 180 | 166 |
| 2006 | 205 | 187 | 196 | 187 | 188 | 179 |
| 2007 | 300 | 294 | 299 | 205 | 282 | 277 |
| 2008 | 335 | 326 | 334 | 289 | 292 | 285 |
| 2009 | 337 | 312 | 336 | 314 | 314 | 290 |
| 2010 | 518 | 501 | 518 | 234 | 235 | 231 |
| 2011 | 409 | 379 | 406 | 333 | 333 | 316 |
| 2012 | 390 | NA | NA | NA | NA | NA |
| 2013 | 283 | NA | NA | NA | NA | NA |
| 2014 | 364 | 349 | 360 | 352 | 353 | 343 |
| Total | 4640 | 3750 | 3936 | 2902 | 3213 | 3089 |

**PART 3 – SITE CLIMATIC INFORMATION**

**Fig S3. Mean daily air temperature in July (A, B), annual temperature norms (C,D) and annual precipitation norms (E, F) across Switzerland in 1961-1990 (A, C, E) and 1981-2010 (B, D, F) (source:** [**www.geo.admin.ch**](http://www.geo.admin.ch)**).** Maps were created by the Federal Office for the Environment (FOEN) and were accessed on the 13.10.23 online at the portal [www.geo.admin.ch](http://www.geo.admin.ch). The legends show how colours correspond to (in each panel from left to right) temperature in graphs A and B, temperature in graphs C and D and rainfall in graphs E and F. The full gradient of colours used in each map is not shown here, but an evenly spaced snapshot of colours across the full gradient is shown. In each graph, the locations of study locations are marked – although these cannot be seen readily in panels B-F.

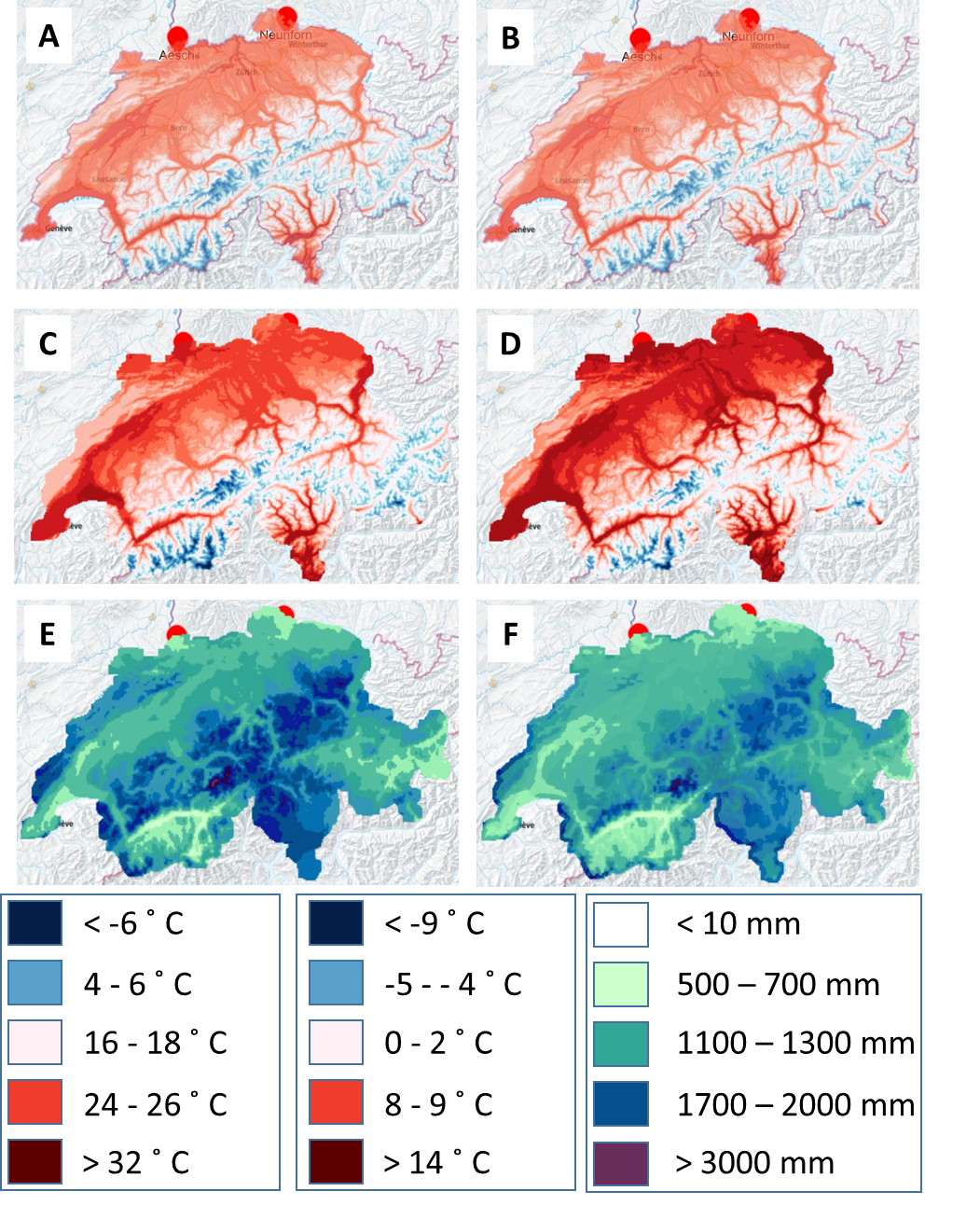

**Fig S4. Annual temperature (A) and precipitation (B) recorded at each site Neunforn (blue-green) and Aesch (red) from 1999-2013.** Temperature measures reflect air temperature 2 m above ground reported as deviation of the annual mean to the norm – the norm being the average air temperature 2m above the ground between 1991 and 2020. Precipitation measures include snowfall, rain and hail and are reported as percentages, and reflect the relation of the annual total to the norm in the period 1991-2020. The dashed lined in each graph shows where points would sit if there were no deviation from the historic norm.

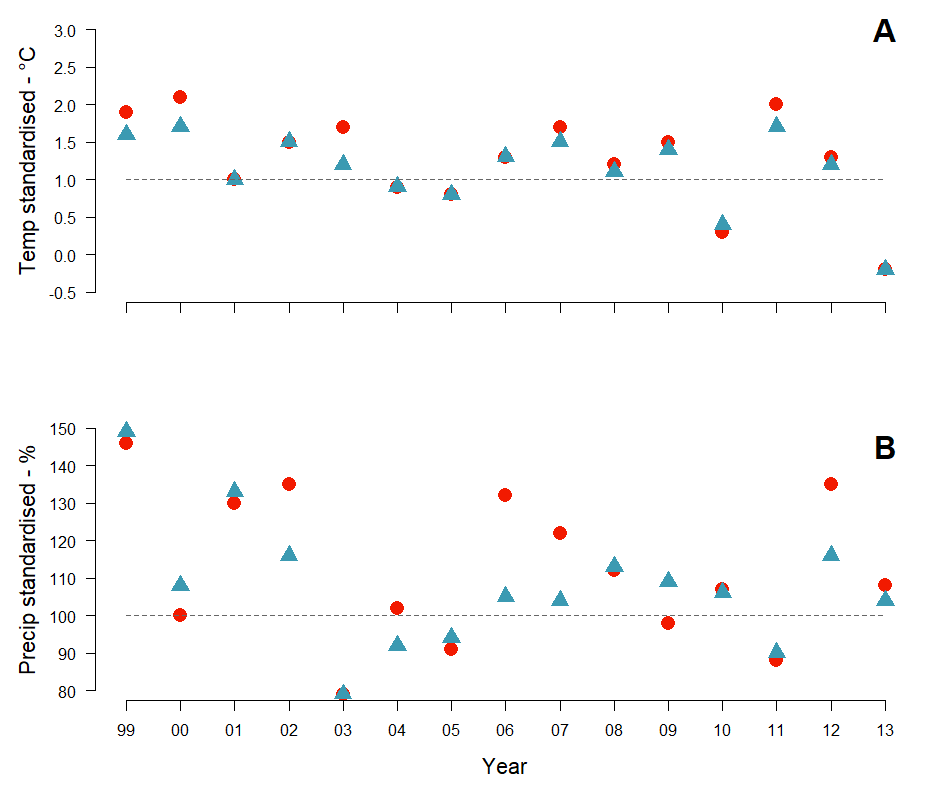

**Fig S5. The correlation between annual temperature and precipitation at Neunforn (blue) and Aesch (red) from 1999-2013.** Regression lines were created using *abline* of *ggplot2*, and show the correlation between climatic variables at each site.

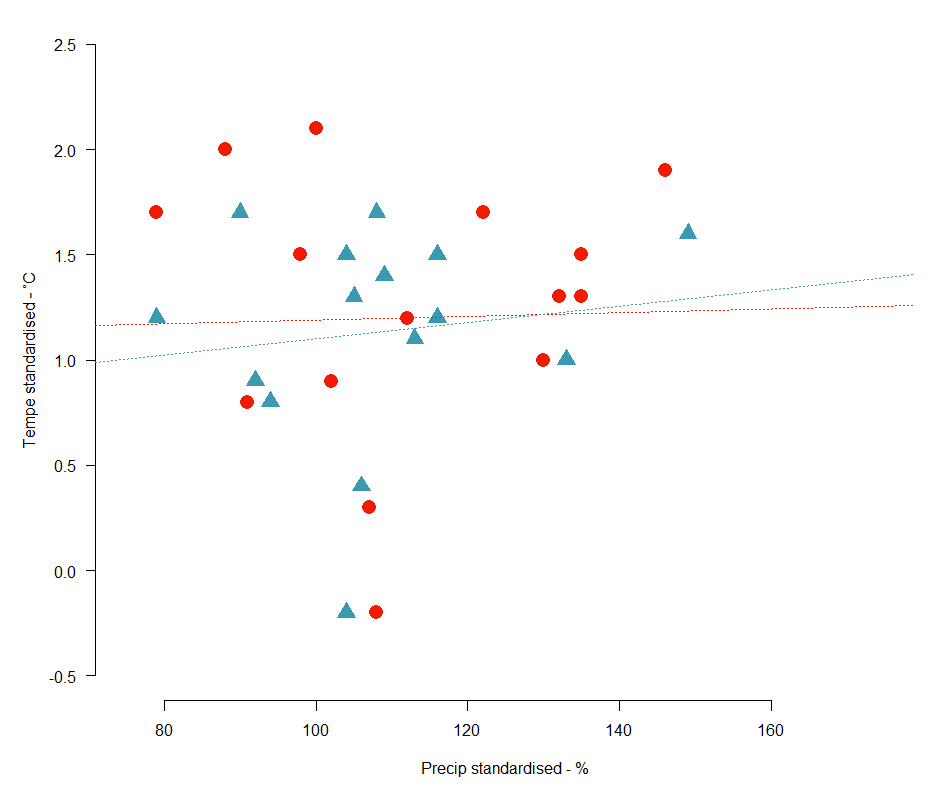

**PART 4 – EXTRA DETAIL STATISTICS**

**Text S1 – Survival Model**

We analyzed survival data in a time-to-event framework, where the event was dying within the observation period (dead = 1, alive = 0) and a “time” is associated with each outcome. For animals that died, this “time” is the approximate age at death (i.e. date of death minus the date each queen was collected from the field). For bees that survived the overall observation period, this age was when bees were recorded as being alive for the last time. These final observations may have been made when noting when queens were transferred to a new box, laid eggs, produced workers or were frozen (e.g. date of egg laying minus the date each queen was collected from the field). Because survival was not monitored at a regular schedule after around mid-June when colonies were established for experiments, all observations are capped on the 166^th^ day of the year. Any bees still alive on this date are recorded as being alive and given a censoring time equivalent to their approximate age on the 166^th^ day of the year (e.g. 166 minus the day of the year each queen was collected from the field). Any survival data that could not be associated with a date was excluded from analyses. Models were analyzed Model structure was otherwise identical to those described for other traits.

**Figure S6. Starting model for the piecewise SEM analyses.** This schematic illustrates the full model used in the piecewise SEM process. Each arrow demonstrates a hypothetical relationship between the variables studied. A) the full model, where climatic variables are presented in dark blue and arrows associated with these variables are solid. Variables associated with where (site) or when (collected DOY) queens were collected are shown in green and associated arrows are shown with narrow dashes. Relationships between response variables are shown as wide dashed arrows linking body mass (purple) with *Crithidia* (grey) and colony establishment (i.e. reproduction – yellow). For clarity, panel B shows only arrows involving climatic variables, panel C shows only collection information and panel D, relationships between response traits. Note that weight was not included as a quadratic term – despite this improving the individual model fit for reproduction. This is because weight is affected by weight as a quadratic term, and this model is assessed as a poor fit.

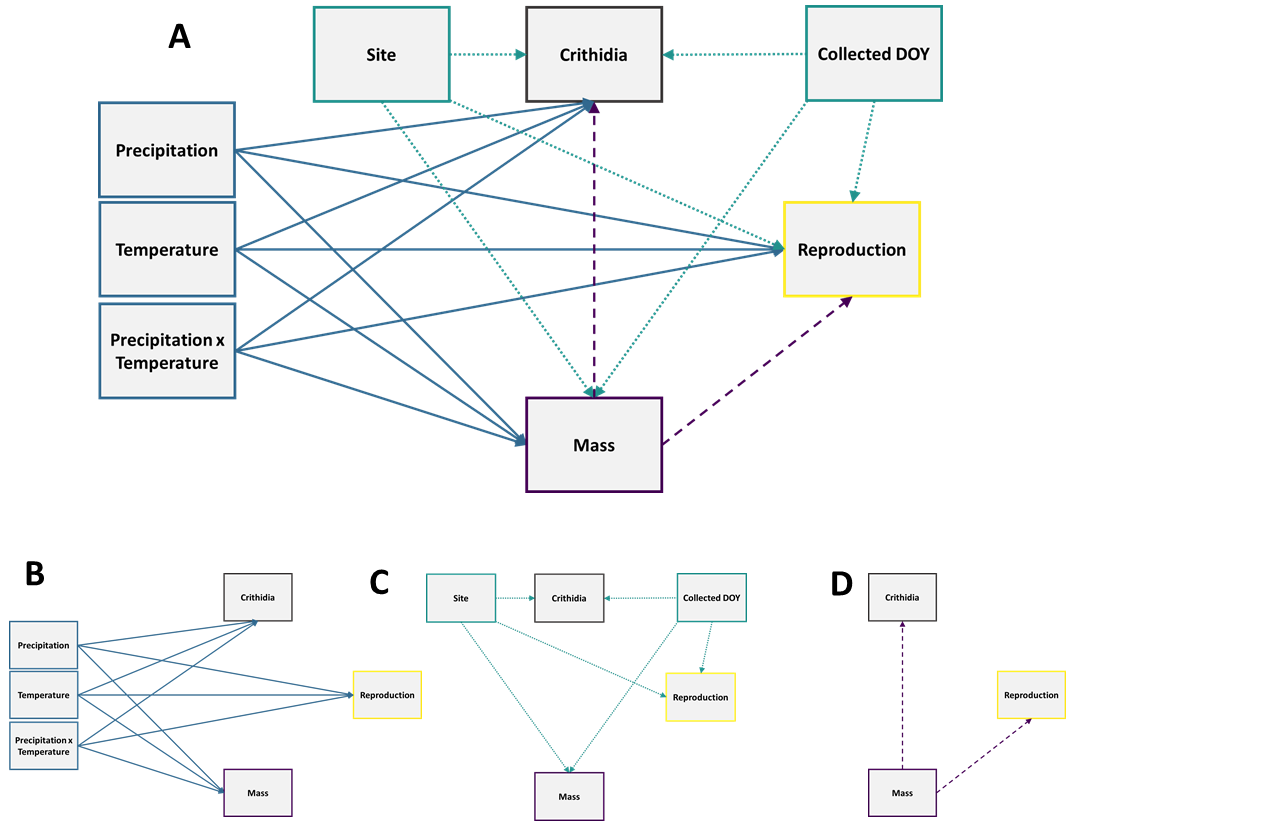

**PART 5 – EXTRA INFORMATION ANNUAL RESULTS**

**Figure S7. The predicted relationship between *Crithidia* infection and queen body mass (g), collection day of the year (DOY) and standardised temperature (ᵒC - linear).** Shaded regions show confidence intervals, and lines show predictions from the preferred model estimated using the *ggpredict* function in the *ggeffects* package. Red = site Aesch, Blue = site Neunforn.

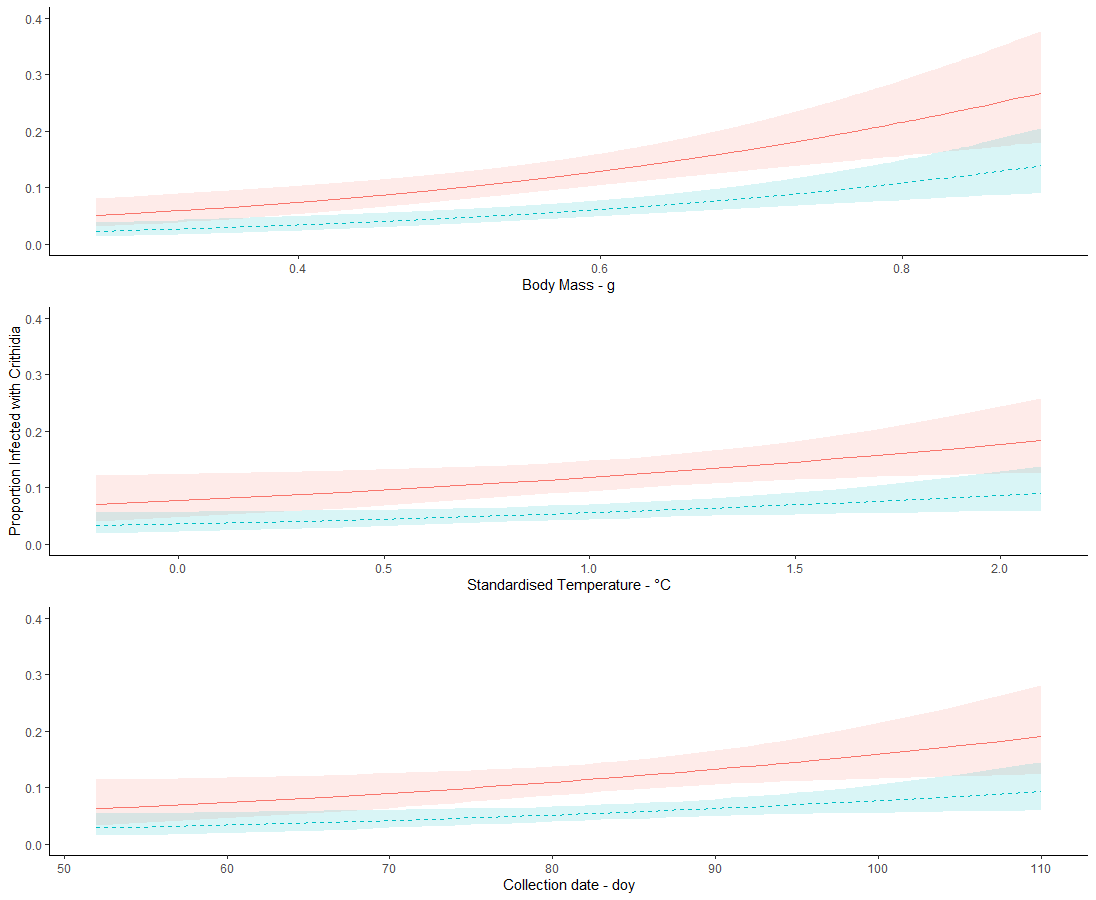

**PART 6 – EXTRA INFORMATION SLIDING WINDOWS**

**Figure S8-11. Output of initial sliding window analyses.** Results are shown separately for each trait but all figures are described here. For each trait there are four graphs – two showing the results for temperature and two for precipitation. For each climatic variable / trait combination, the “model fit” graph show the difference in AIC values between a model where a window opens in each time step, and closes in each time step, relative to a model that excludes climate. The greater divergence between these values, the better the fit. Colour key (with ∆AIC values) is indicated in the graph: red values show large ∆AIC values, showing that a model performs well compared to the baseline model, while purple values show a minimal difference between the baseline model and climate model. The graphs describing “effect” summarise “beta” and show the relationship between climate and trait at each point. The resolution of the graphs for temperature and precipitation differ – results for temperature show weekly steps, precipitation monthly steps. Window 0 = 1^st^ March in year queens collected.

**Fig S8. Sliding Window Analyses Results Summary – Body Mass**

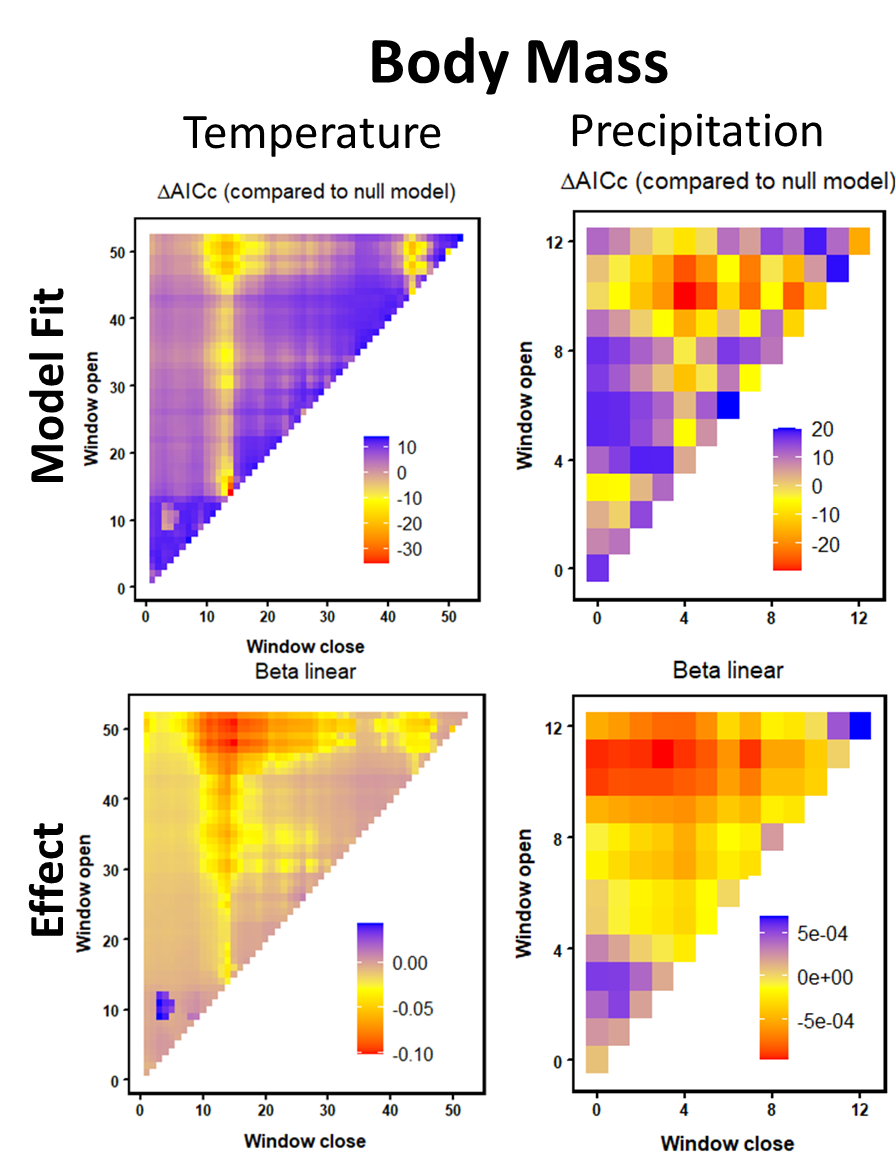

**Fig S9. Sliding Window Analyses Results Summary – *Crithidia***

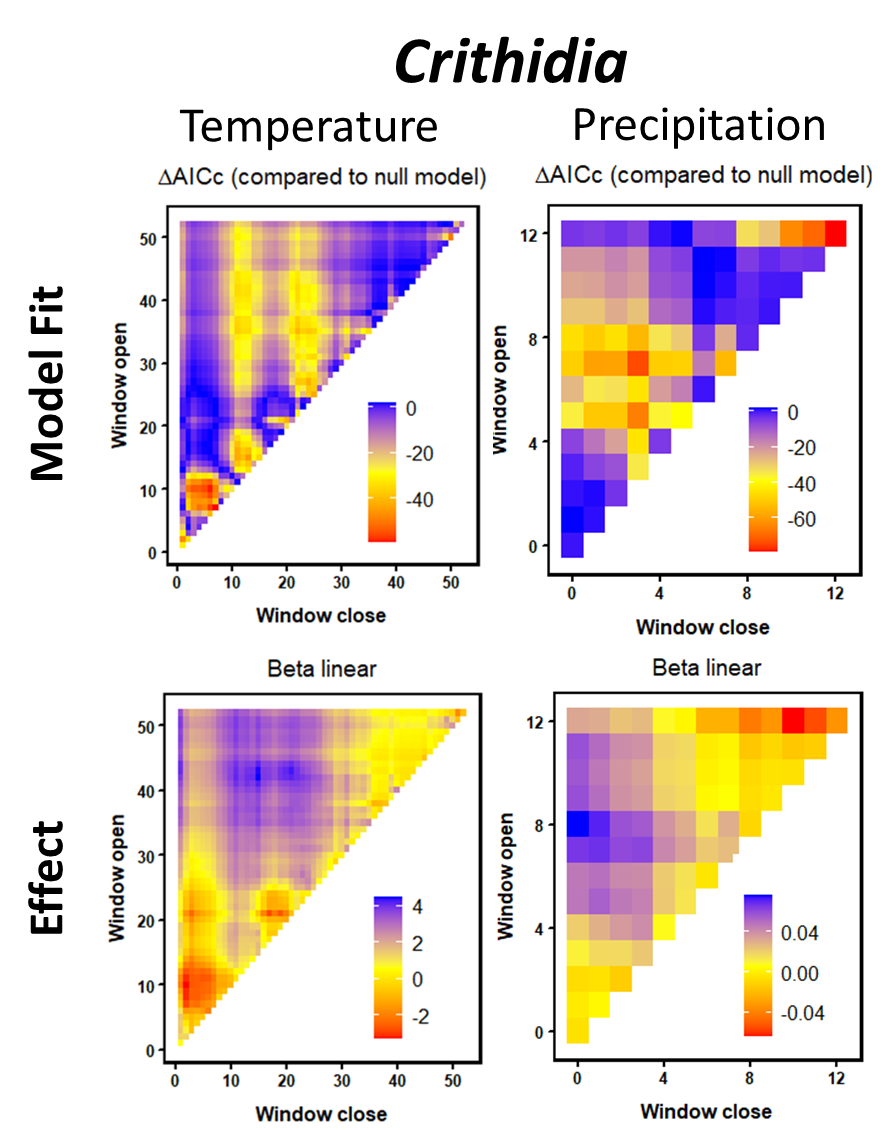

**Fig S10. Sliding Window Analyses Results Summary – Colony Establishment i.e. Reproduction.**

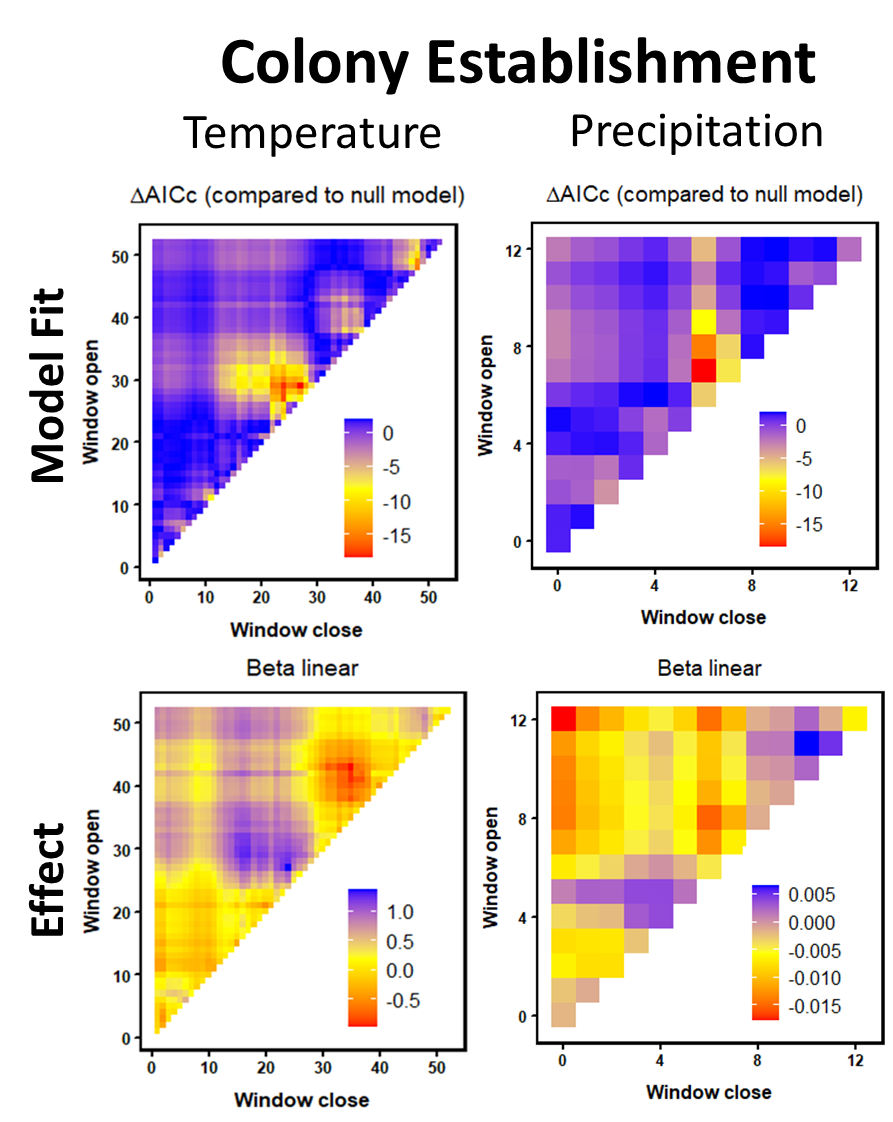

**Fig S11. Sliding Window Analyses Results Summary – Survival**

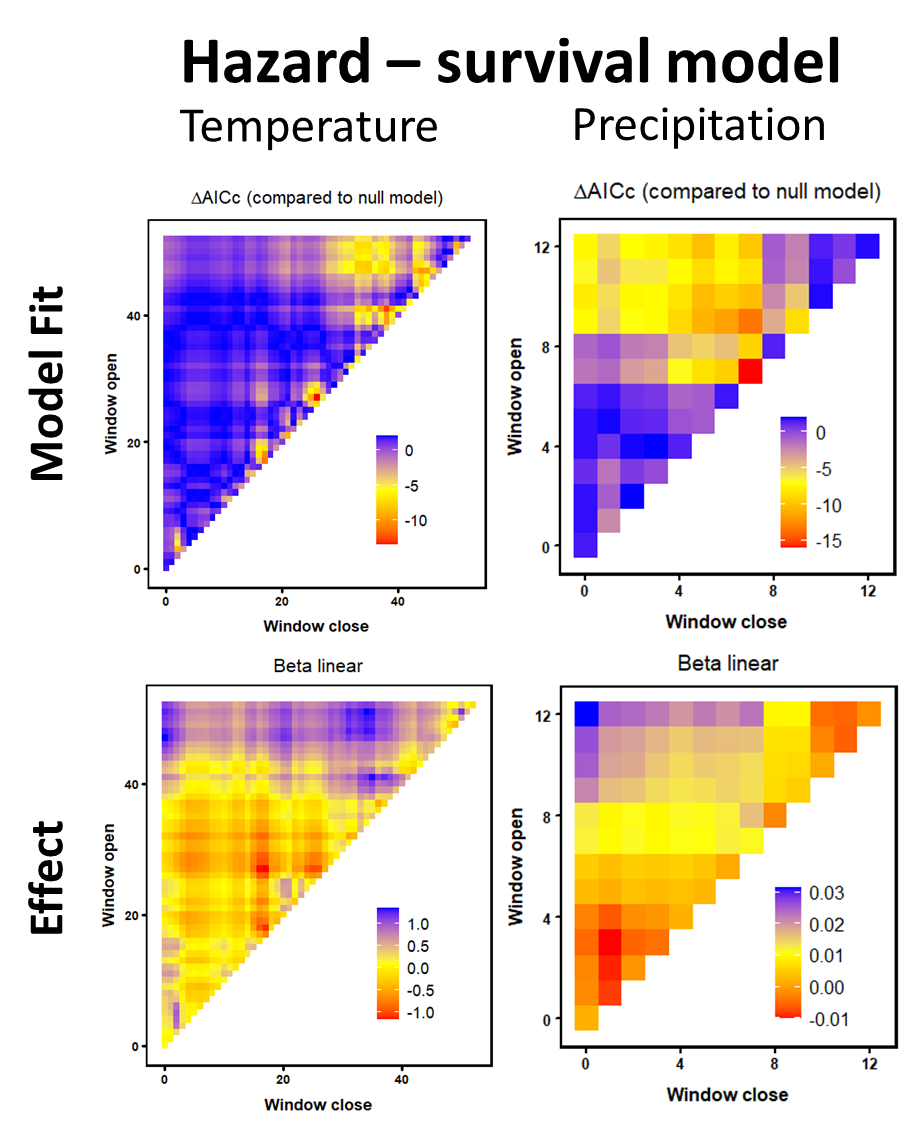

**Table S3. Dates that each window in sliding window analyses corresponds.** These dates are for a non-leap year.

| **Window – Temp** | **Corresponding Date** | **Window - Precipitation** |
| --- | --- | --- |
| 0 | 1^st^ March | 0 |
| 1 | 22^nd^ Feb | 1 |
| 2 | 15^th^ Feb |  |
| 3 | 8^th^ Feb |  |
| 4 | 1^st^ Feb |  |
| 5 | 25^th^ Jan | 2 |
| 6 | 18^th^ Jan |  |
| 7 | 11^th^ Jan |  |
| 8 | 4^th^ Jan |  |
| 9 | 28^th^ Dec | 2 |
| 10 | 21^st^ Dec |  |
| 11 | 14^th^ Dec |  |
| 12 | 7^th^ Dec |  |
| 13 | 30^th^ Nov | 4 |
| 14 | 23^rd^ Nov |  |
| 15 | 16^th^ Nov |  |
| 16 | 9^th^ Nov |  |
| 17 | 2^nd^ Nov |  |
| 18 | 26^th^ Oct | 5 |
| 19 | 19^th^ Oct |  |
| 20 | 12^th^ Oct |  |
| 21 | 5^th^ Oct |  |
| 22 | 28^th^ Sept | 6 |
| 23 | 21^st^ Sept |  |
| 24 | 14^th^ Sept |  |
| 25 | 7^th^ Sept |  |
| 26 | 31^st^ Aug | 7 |
| 27 | 24^th^ Aug |  |
| 28 | 17^th^ Aug |  |
| 29 | 10^th^ Aug |  |
| 30 | 3^rd^ Aug |  |
| 31 | 27^th^ July | 8 |
| 32 | 20^th^ July |  |
| 33 | 13^th^ July |  |
| 34 | 6^th^ July |  |
| 35 | 29^th^ June | 9 |
| 36 | 22^nd^ June |  |
| 37 | 15^th^ June |  |
| 38 | 8^th^ June |  |
| 39 | 1^st^ June |  |
| 40 | 25^th^ May | 10 |
| 41 | 18^th^ May |  |
| 42 | 11^th^ May |  |
| 43 | 4^th^ May |  |
| 44 | 27^th^ April | 11 |
| 45 | 20^th^ April |  |
| 46 | 13^th^ April |  |
| 47 | 6^th^ April |  |
| 48 | 30^th^ March | 12 |
| 49 | 23^rd^ March |  |
| 50 | 16^th^ March |  |
| 51 | 9^th^ March |  |
| 52 | 2^nd^ March |  |

**Table S4. Results of moving window analyses for (A) Precipitation and (B) Temperature data.**

| **Model** | **Window** | | **∆AIC** | ***ß*** | **SE** | ***Wi*** | ***P ΔAICc*** | ***P*_C_** |
| --- | --- | --- | --- | --- | --- | --- | --- | --- |
|  | **Start** | **End** |  |  |  |  |  |  |
| ***Precipitation*** |  |  |  |  |  |  |  |  |
| ***Body Mass (g)*** |  |  |  |  |  |  |  |  |
| *Best Model*  *Confidence Set* | 10  (10) | 4  (4.5) | -29.45 | -0.001  (-0.001) | 0.003 | 0.674 | **< 0.001** | **0.009** |
| ***Crithidia* (0 / 1)** |  |  |  |  |  |  |  |  |
| *Best Model* | 12 | 12 | -78.94 | -0.034 | 0.058 | 0.926 | **< 0.001** | **< 0.001** |
| **Reproduction (0 / 1)** |  |  |  |  |  |  |  |  |
| *Best Model* | 7 | 6 | -18.46 | -0.013 | 0.038 | 0.826 | **< 0.001** | **< 0.001** |
| **Mortality risk** |  |  |  |  |  |  |  |  |
| *Best Model*  *Confidence Set* | 7  (10) | 7  (5) | -15.93 | 0.011  (0.013) | 0.003 | 0.464 | **< 0.001** | **0.007** |
| ***Temperature*** |  |  |  |  |  |  |  |  |
| **Mass (g)** |  |  |  |  |  |  |  |  |
| *Best Model* | 14 | 14 | -35.87 | -0.025 | 0.003 | 0.920 | **< 0.001** | **0.002** |
| ***Crithidia*** |  |  |  |  |  |  |  |  |
| *Best Model*  *Confidence Set* | 7  (10) | 7  (6) | -59.18 | -1.237  (-1.709) | 0.060 | 0.377 | **< 0.001** | **0.001** |
| **Reproduction (0 / 1)** |  |  |  |  |  |  |  |  |
| *Best Model*  *Confidence Set* | 29  (30) | 27 (23) | -18.41 | 0.972  (1.020) | 0.039 | 0.174 | **< 0.001** | **0.001** |
| ***Mortality risk*** |  |  |  |  |  |  |  |  |
| Range – all year |  |  |  |  |  |  |  |  |
| *Best Model* | 27 | 26 | -13.50 | -0.992 | 0.257 | 0.132 |  |  |
| Range 1-19 |  |  |  |  |  |  |  |  |
| *Best Model*  *Confidence Set* | 17  (13) | 17  (6) | -11.30 | -0.714  (-0.496) | 0.201 | 0.368 | **< 0.001** | **0.036** |
| Range 20-29 |  |  |  |  |  |  |  |  |
| *Best Model*  *Confidence Set* | 27  (27) | 26  (25.5) | -13.50 | -0.992  (-0.806) | 0.257 | 0.691 | **< 0.001** | **0.003** |
| Range 30-52 |  |  |  |  |  |  |  |  |
| *Best Model*  *Confidence Set* | 41  (48) | 38  (36) | -12.64 | 1.089  (0.904) | 0.285 | 0.147 | **0.002** | **0.045** |

Windows for precipitation data are measured in months, while windows for temperature data are measured in weeks. For traits where there was one clear climate peak, and the best model from the sliding window analyses had a high probability of being the best model in the model set, we only report parameters associated with the best model. For traits where there was one clear climate peak, but multiple models have a similar probability of being the best model in the model set, we also report a *ß* value based on model averaging on the confidence set - a subset of models where we can be 95% confident that the best model is included – and median windows for the set. These values are provided in parenthesis. For the models relating climate to survival, the three best fitting models each corresponded to three distinct peaks. Thus, individual moving windows were run for periods corresponding to each peak (1 – 19 weeks prior to 1.3; 20 – 29 weeks, 30 – 52 weeks). The confidence set was calculated for each of these ranges and significance testing carried out for each peak. **∆AIC** - the AIC value of the model reported, subtracted from the AIC of the baseline model (i.e. a model including climate only). These values are comparable across traits – more negative values indicate a stronger influence of the climatic variable tested in that model. ***ß –*** relationship between the climatic variable tested and the response variable. For example, the -0.001 *ß* for the model linking body mass and precipitation, indicates that for each increase in precipitation (% relative to historical norm), body mass declines by -0.001g. Note that because survival data were analysed in a cox model framework, here beta reflects the reduction in hazard i.e. risk of dying. **SE** – standard error around the estimate of *ß.* ***Wi –*** the probability that the model reported, is the best model within the model set. ***PΔAICc, P*_C_ *–*** results of randomisation analyses, the likelihood of obtaining the level of model support observed for the best model by chance. These values were always calculated with 500 iterations and degrees of freedom calculated as the number of years of data included in the analyses multiplied by the number of sites. All models remain significant even if using half as many degrees of freedom (i.e. based on how many years of data are included only).
